# Chromosomal integration of *bla*_CTX-M_ genes in diverse *Escherichia coli* isolates recovered from river water in Japan

**DOI:** 10.1101/2022.03.05.483147

**Authors:** Ryota Gomi, Masaki Yamamoto, Michio Tanaka, Yasufumi Matsumura

## Abstract

Occurrence of extended-spectrum β-lactamase (ESBL)-producing *Escherichia coli* (ESBLEC) in environmental waters is of great concern. However, unlike clinical ESBLEC, their genetic characteristics, in particular the genetic contexts of ESBL genes, are not well understood. In this study, we sequenced and analyzed the genomes of CTX-M-producing *E. coli* isolates recovered from river water to fully characterize the genetic contexts of *bla*_CTX-M_ genes. Among the 14 isolates with completed genomes, *bla*_CTX-M_ genes were detected on the chromosome in nine isolates. All but one chromosomal *bla*_CTX-M_ genes were associated with IS*Ecp1* and were carried on different transposition units ranging in size from 2,855 bp to 11,093 bp; the exception, *bla*_CTX-M-2_, was associated with IS*CR1*. The remaining five isolates carried *bla*_CTX-M_ genes on epidemic IncI1 plasmids of different sequence types (STs) (ST3, ST16, ST113, and ST167) (n = 4) or on an IncB/O/K/Z plasmid (n = 1). This study revealed that environmental *E. coli* carry *bla*_CTX-M_ genes in diverse genetic contexts. Apparent high prevalence of chromosomal *bla*_CTX-M_ potentially indicates that some *E. coli* can stably maintain *bla*_CTX-M_ genes in environmental waters, though further studies are needed to confirm this.

## 1. Introduction

Occurrence of antibiotic-resistant bacteria (ARB) in surface waters is a public health concern. Among ARB, extended-spectrum-β-lactamase (ESBL)-producing *Escherichia coli* (ESBLEC) represent a major threat to public health, because ESBLEC can be important causes of community-onset infections for which treatment options are limited (Pitout, 2010; Pitout and Laupland, 2008). ESBL enzymes are divided into several families (e.g., CTX-M and ESBL variants of TEM and SHV), among which CTX-M enzymes have been recognized as the most common ESBL type (Castanheira et al., 2021). Previous studies have reported the predominance of CTX-M ESBLs among environmental *E. coli*. For example, CTX-M-14(-like) was prevalent among ESBLEC in surface waters in China, Japan, and Korea, and CTX-M-15 was prevalent among ESBLEC in surface waters in France, Germany, and Switzerland (Falgenhauer et al., 2021; Girlich et al., 2020; Jang et al., 2013; Liu et al., 2018; Miyagi and Hirai, 2019; Zurfluh et al., 2013).

ESBL genes are often found on plasmids, which facilitates their horizontal spread (Castanheira et al., 2021). However, some previous studies also detected ESBL genes on chromosomes and suggested that integration of ESBL genes into chromosomes could contribute to the stabilization and maintenance of ESBL genes in the absence of antibiotic selection pressure (Hirai et al., 2013; Nair et al., 2021; Rodriguez et al., 2014). Thus, information on the location (chromosome or plasmid) of ESBL genes is important to predict the mobility and stability of these genes. Although, previous studies have reported the occurrence of ESBLEC in environmental waters, limited information exists regarding the locations and genetic contexts of ESBL genes in environmental ESBLEC (Falgenhauer et al., 2021; Girlich et al., 2020; Liu et al., 2018). Here, we completely sequenced the genomes of CTX-M-producing *E. coli* isolates recovered from river water in Japan, aiming to elucidate the locations and genetic contexts of the *bla*_CTX-M_ genes in environmental ESBLEC. The genetic contexts of chromosomal *bla*_CTX-M_ genes in the genomes of environmental *E. coli* retrieved from the NCBI Reference Sequence Database (RefSeq) were also characterized.

## 2. Materials and methods

### 2.1. Bacterial isolates

We previously collected 531 *E. coli* isolates from the Yamato River in Japan between 2011 and 2013 and detected 18 isolates with *bla*_CTX-M_ genes (Gomi et al., 2014; Gomi et al., 2017). Briefly, we collected a total of 27 water samples from 10 sites in the Yamato River in August and November of 2011, February of 2012, and October of 2013 (note that water samples were not collected from all sampling sites on each sampling occasion). The samples were processed using the membrane filter method with XM-G agar (Nissui, Tokyo, Japan) to obtain *E. coli* isolates. In total, 531 *E. coli* isolates were obtained, and a subset of isolates (n = 155) was selected and subjected to Illumina short-read sequencing. Among the sequenced isolates, we detected 18 isolates that carried *bla*_CTX-M_ genes. These 18 isolates were analyzed as described below.

### 2.2. Antibiotic susceptibility testing

Antibiotic susceptibility was assessed by microdilution using the dry plate Eiken assay (Eiken, Tokyo, Japan) as described previously (Gomi et al., 2017) (see **Table S1** for the antimicrobial agents used). The results were interpreted according to CLSI criteria (CLSI, 2020) and EUCAST epidemiological cutoff (ECOFF) values (http://www.eucast.org). ESBL production was confirmed following the CLSI guidelines (CLSI, 2020). A detailed description of antibiotic susceptibility testing is provided in the Supplementary Materials and methods.

### 2.3. Genome sequencing and assembly

DNA was extracted from each isolate using a DNeasy PowerSoil Pro Kit (Qiagen, Hilden, Germany). The short-read sequencing library was prepared using the Illumina DNA Prep (Illumina, San Diego, CA), and the library was sequenced on an Illumina NovaSeq 6000 system with 250-bp paired-end sequencing (although we obtained Illumina reads for the isolates in our previous study (Gomi et al., 2017), the Illumina reads had low depth/uneven coverage and hence we sequenced the isolates again with Illumina technology in the present study). The long-read sequencing library was prepared using the SQK-LSK109 kit (Oxford Nanopore Technologies, Oxford, UK), and the prepared library was sequenced on the MinION with a FLO-MIN106 flow cell.

Illumina short reads were subsampled using seqtk (v1.3, https://github.com/lh3/seqtk), and the subsampled reads were trimmed with fastp (v0.20.0) (Chen et al., 2018). Oxford Nanopore Technologies (ONT) long reads were filtered with Filtlong (v0.2.0, https://github.com/rrwick/Filtlong) to remove reads with low quality or short length. Hybrid assembly of Illumina short reads and ONT long reads was performed by employing a short-read-first approach, i.e., Unicycler (v0.4.8) with the --no_correct option (Wick et al., 2017). Hybrid assembly was also performed using a long-read-first approach, i.e., long-read assembly by Flye (v2.9) (Kolmogorov et al., 2019) followed by long-read polishing using Medaka (v1.4.4, https://github.com/nanoporetech/medaka) and short-read polishing using Polypolish (v0.4.3) (Wick and Holt, 2022). Assembly graphs of both hybrid assemblies were visualized using Bandage (v0.8.1) for each isolate (Wick et al., 2015), and a better assembly was chosen as the final assembly.

### 2.4. Genomic analysis

Genomes were annotated using the RAST server (Aziz et al., 2008), ISfinder (Siguier et al., 2006), ResFinder 4.1 (Bortolaia et al., 2020), and the BLAST algorithm (https://blast.ncbi.nlm.nih.gov/Blast.cgi). Plasmid replicons were detected using PlasmidFinder 2.1 (Carattoli et al., 2014). Relaxase types, mate-pair formation types, and transferability of plasmids were determined using MOB-typer (Robertson and Nash, 2018). IncI1 plasmids were annotated using plasmids R64, ColIb-P9, and R621a as references (Carattoli et al., 2021).

### 2.5. Characterization of environmental *E. coli* genomes with *bla*_CTX-M_ genes in RefSeq

RefSeq *E. coli* genomes with an assembly level of “complete genome” (n = 1903) were downloaded in January 2022. Metadata information, such as collection date and isolation source, was extracted from the GenBank files using an in-house Python script, and genomes of *E. coli* from environmental waters were retrieved (n = 53). Antibiotic resistance genes were detected using ABRicate (https://github.com/tseemann/abricate), and genomes carrying *bla*_CTX-M_ on the chromosome were analyzed as described above.

### 2.6. Accession numbers

The complete genomes and sequence read files were deposited in GenBank and the NCBI SRA under BioProject PRJNA800231 (also see **Table S2** for the accession number of each replicon for the completed genomes and **Table S1** for SRA accession numbers of the remaining genomes).

## 3. Results and discussion

### 3.1. Basic characteristics of *E. coli* isolates sequenced in this study

All 18 *bla*_CTX-M_-carrying isolates characterized in the present study were non-susceptible/non-wild type to ampicillin, aztreonam, cefazolin, cefepime, cefotaxime, and cefpodoxime, while all were susceptible/wild type to amoxicillin-clavulanic acid, piperacillin-tazobactam, imipenem, meropenem, amikacin, and colistin. Non-susceptibility/non-wild type rates for the remaining antibiotics ranged from 6% (fosfomycin) to 89% (ampicillin-sulbactam) (**Table S1**). ESBL production was confirmed in all the isolates. The 18 isolates were highly diverse, comprising 16 different sequence types (STs), namely ST38 (n = 2), ST69 (n = 2), and a single isolate from each of ST10, ST46, ST117, ST216, ST354, ST457, ST744, ST1193, ST2003, ST3107, ST6214, ST6215, ST6216, and ST6220 (**Table S1**). The *bla*_CTX-M_ genes detected were also diverse, namely *bla*_CTX-M-14_ (n = 8), *bla*_CTX-M-2_ (n = 2), *bla*_CTX-M-15_ (n = 2), *bla*_CTX-M-24_ (n = 2), *bla*_CTX-M-55_ (n = 2), *bla*_CTX-M-1_ (n = 1), *bla*_CTX-M-3_ (n = 1), and *bla*_CTX-M-8_ (n = 1). One isolate (KTa008) carried two copies of the *bla*_CTX-M-14_ gene. In Japan, ESBLEC are frequently detected among *E. coli* infections. For example, 22% of extraintestinal pathogenic *E. coli* isolates were positive for one or more ESBL genes in a study conducted at 10 acute-care hospitals (Matsumura et al., 2017a). Like in other countries, these ESBLEC infections in Japan have been shown to be mainly due to CTX-M-producing ST131 (Matsumura et al., 2017b). However, the ESBLEC isolates analyzed in the present study belonged to a wide variety of STs and none belonged to ST131, which indicates that the clonal composition of ESBLEC between environmental settings and clinical settings may be different in Japan.

Of the 18 isolates, the genomes of 14 could be completed. These genomes carried 0 to 8 plasmids (see **Table S2** for details). The genomes of the remaining four isolates could not be completed, probably due to the presence of long repeats in the genomes. The location and genetic contexts of the *bla*_CTX-M_ genes were determined for the 14 isolates with completed genomes. Of these 14, nine carried *bla*_CTX-M_ genes on chromosomes and five carried *bla*_CTX-M_ genes on plasmids.

### 3.2. Genetic contexts of chromosomal *bla*_CTX-M_ genes

In eight of the nine isolates which carried *bla*_CTX-M_ genes on chromosomes, the *bla*_CTX-M_ genes were associated with IS*Ecp1* and carried on transposition units (TPU) of different sizes (2,855 bp to 11,093 bp) (**Figure 1**). These TPU were flanked by AT-rich 5-bp target site duplications (TSD). None of the TPU were identical, and the insertion sites of the TPU were different in all cases, indicating that different transposition events contributed to the chromosomal integration of each TPU. Notably, the TPU in KFu023 contained multiple resistance genes, including *bla*_CTX-M-15_, *aac(6′)-Ib-cr, bla*_OXA-1_, and *ΔcatB3*. This seems to highlight the ability of IS*Ecp1* to capture sequences of different lengths in different transposition events and thus simultaneously mobilize antibiotic resistance genes of different origins (Partridge et al., 2018). In most cases, these TPU were directly inserted into chromosomes. However, in KKa019, the TPU was part of a 35,625 bp multiresistance region bracketed by two copies of IS*26* (annotated as IS*15DI* in the ISFinder database), and the whole multiresistance region was flanked by 8-bp TSD (CGTTGCCG) in the chromosome (**Figure 2a**). KTa008 carried two copies of *bla*_CTX-M-14_, which were separated by ∼1,169 kbp in the chromosome. Analysis of the genetic contexts of these *bla*_CTX-M-14_ genes indicated transposition of an already chromosomally-located TPU and an adjacent region into another chromosomal location (**Figure S1**). This type of ‘recurrent’ transposition was previously reported in clinical ESBLEC, though the TPU and insertion sites were different (Hamamoto and Hirai, 2019).

**Figure 1.**
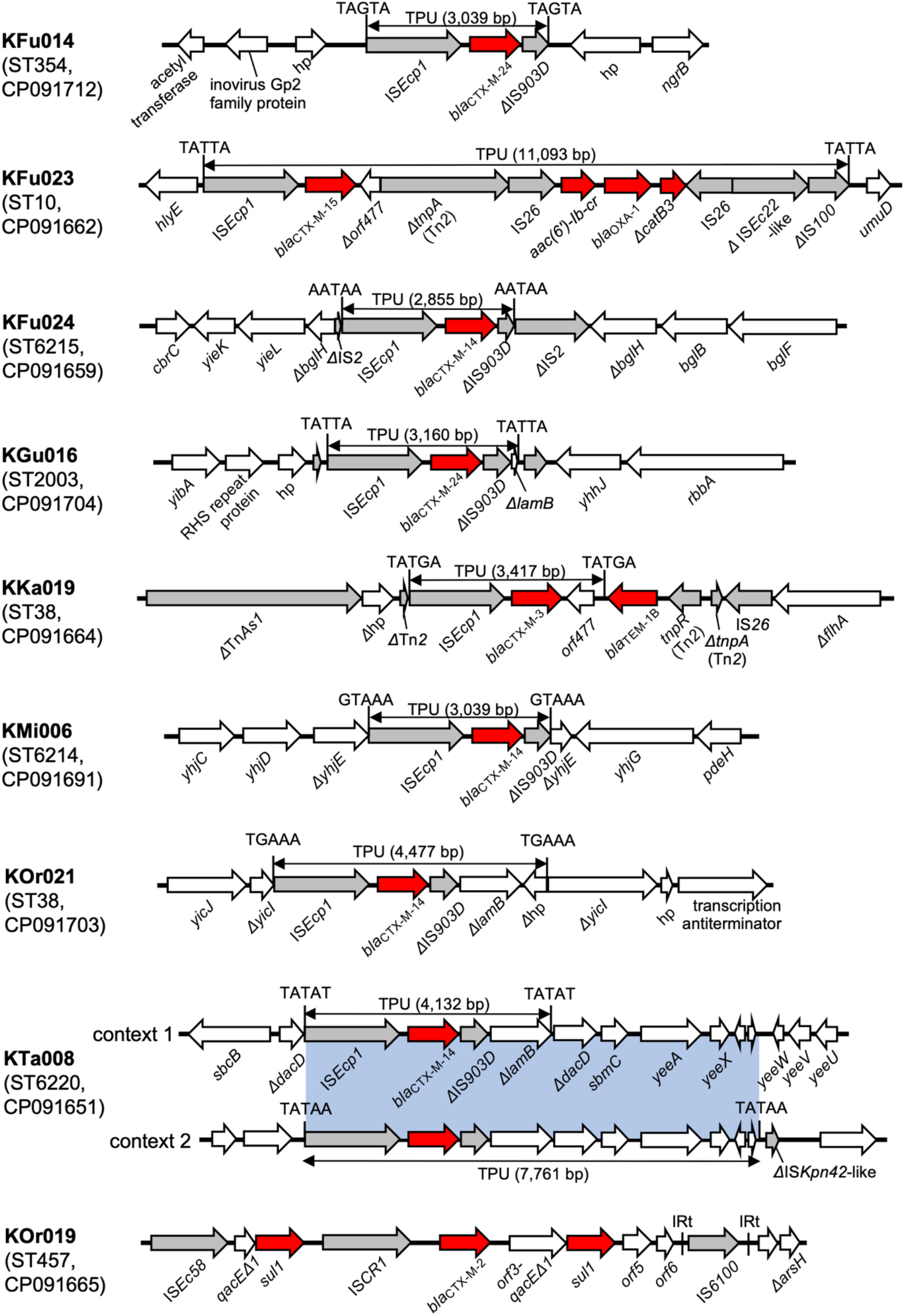
Genetic contexts of chromosomal *bla*_CTX-M_ genes. Red arrows indicate antibiotic resistance genes, gray arrows indicate mobile elements, and white arrows indicate other genes. The blue shaded box indicates the regions with 100% nucleotide identity. The entire resistance regions of KKa019 and KOr019 are shown in **Figure 2**. 5-bp TSD are shown next to each TPU. hp: hypothetical protein.

**Figure 2.**
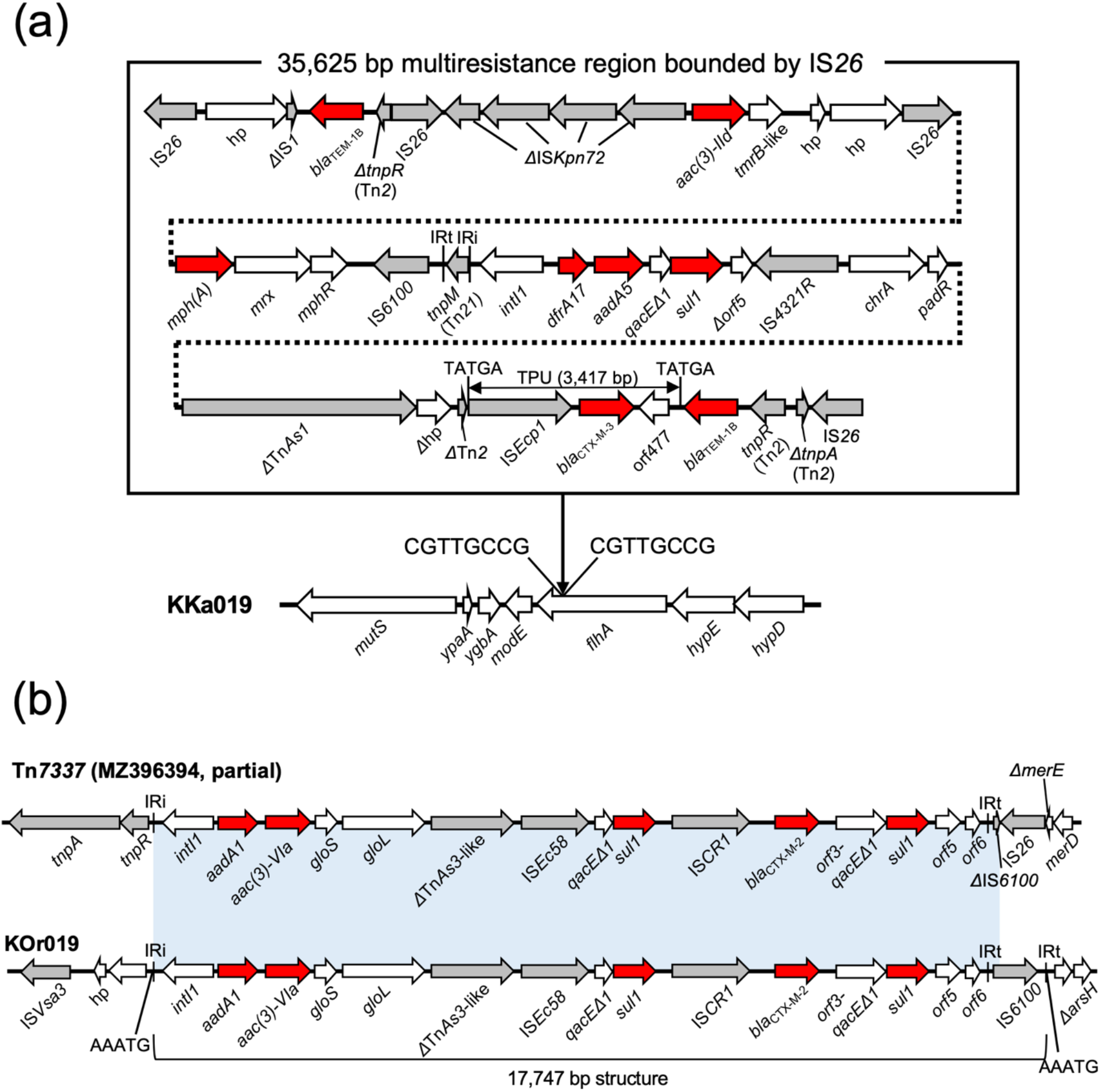
(a) A multiresistance region on the chromosome of KKa019. Insertion of the resistance region into *flhA* generated 8-bp TSD of CGTTGCCG. A BLASTN search against the nucleotide collection (nr/nt) database using the 35,625 bp multiresistance region as a query sequence identified some related structures, but none showed >90% coverage. (b) A multiresistance region on the chromosome of KOr019. Tn*7337* is shown for the purpose of comparison. The 17,747 bp structure from IRi to IRt was inserted into an intergenic region in the KOr019 chromosome, generating 5-bp TSD of AAATG. The light blue shaded box indicates regions with >99% nucleotide identity.

The above results indicate that analysis of sequences immediately upstream and downstream of *bla*_CTX-M_ (e.g., identification of IS*Ecp1* upstream and *orf477* downstream of *bla*_CTX-M_) is not sufficient to track the movement and evolution of *bla*_CTX-M_, since TPU can be very long (sometimes > 10,000 bp) or can be inserted into a large multiresistance region. This can be overcome, for example, by determining the complete genomes as shown in the present study. We searched for *E. coli* genomes in the public database which carry the same TPU in the same chromosomal location as those identified in the present study. BLASTN searches were performed against the nucleotide collection (nr/nt) database in January 2022 using the TPU and surrounding sequences identified in the present study as query sequences. Three of the nine TPU/insertion site combinations were found in the database, while the remaining six were not found, potentially corresponding to novel integration events (**Table S3**).

In KOr019, *bla*_CTX-M-2_ was associated with IS*CR1* and situated within a complex class 1 integron, which is congruent with previous studies (Canton et al., 2012; Partridge et al., 2018). This complex class 1 integron was closely related to a recently reported transposon Tn*7337* (**Figure 2b**) (Coppola et al., 2022). KOr019 carried the IRt (inverted repeat at *tni* end)-IS*6100*-IRt structure, but this structure was truncated by IS*26* in Tn*7337*. The 17,747 bp structure from IRi (inverted repeat at *intI*1 end) to IRt was inserted into the chromosome of KOr019, creating 5-bp TSD of AAATG. This insertion seems to have been mediated by Tni proteins provided *in trans*, since the *tni* genes in the complex class 1 integron were absent (Partridge et al., 2018).

### 3.3. Genetic contexts of *bla*_CTX-M_ genes on plasmids

The five completed plasmids with *bla*_CTX-M_ genes belonged to IncI1 (n = 4) and IncB/O/K/Z (n = 1) (**Figure 3a**). IncI1 plasmids were typed as four different STs, namely ST3, ST16, ST113, and ST167, by IncI1 pMLST (Carattoli et al., 2014). TPU were inserted into plasmid backbones in the IncI1/ST3, IncI1/ST16, and IncI1/ST167 plasmids. Interestingly, the IncI1/ST16 and IncI1/ST167 plasmids carried the same TPU (except for insertion of IS*1* in the IncI1/ST16 plasmid) inserted into the same location. IncI1/ST167 is a single-locus variant of IncI1/ST16, which indicates the TPU was inserted into the plasmid backbone before the ST diversification. In the IncI1/ST113 plasmid, an IS26-ΔIS*10R*-*bla*_CTX-M-8_-IS*26* structure was inserted into the plasmid. BLASTN analysis identified IncI1 plasmids belonging to the same IncI1-STs and carrying the same TPU/IS*26* structures inserted into the same locations from different countries (e.g., MH847453 from France for *bla*_CTX-M-1_-carrrying IncI1/ST3, CP056650 from the UK for *bla*_CTX-M-55_-carrrying IncI1/ST16, KY964068 from Portugal for *bla*_CTX-M-8_-carrrying IncI1/ST113, CP021208 from China for *bla*_CTX-M-55_-carrrying IncI1/ST167), which indicates circulation of epidemic CTX-M IncI1 plasmids in geographically distant countries.

**Figure 3.**
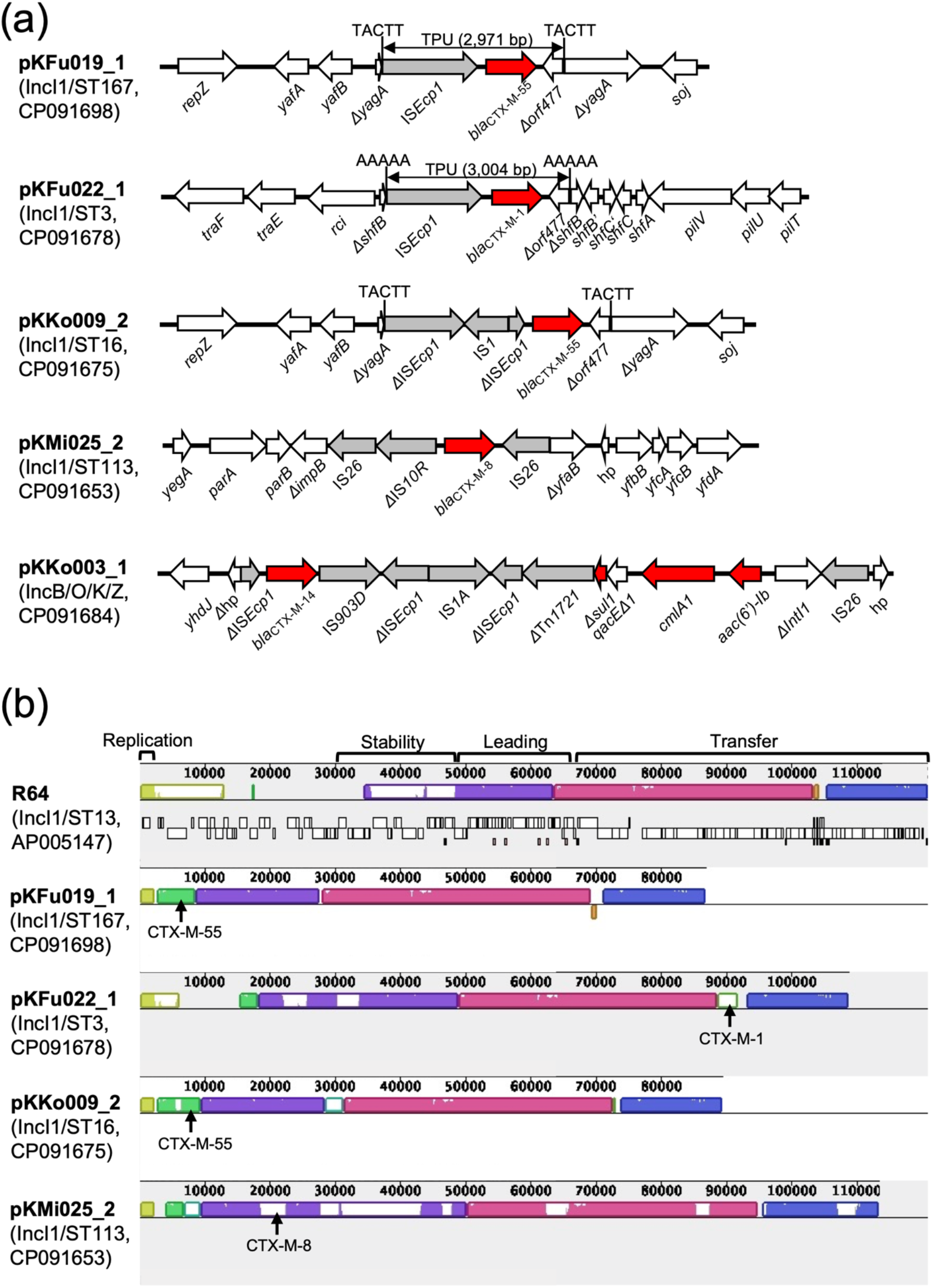
(a) Genetic contexts of *bla*_CTX-M_ genes on plasmids. (b) Mauve comparison of IncI1 plasmids (Darling et al., 2010). IncI1 reference plasmid R64 is shown at the top. Four major regions in R64 (replication, stability, leading, and conjugative transfer) are indicated. Homologous segments are shown as colored blocks. Locations of *bla*_CTX-M_ genes are indicated with arrows.

Comparison of IncI1 plasmids revealed the replication region, the leading region, and the transfer regions in the IncI1 reference plasmid R64 were well conserved in all IncI1 plasmids detected in this study, while many genes in the stability region were lacking (**Figure 3b**). In the IncB/O/K/Z plasmid, *bla*_CTX-M-14_ was linked to ΔTn*1721* and a class 1 integron. Similar genetic contexts were previously reported (e.g., JF701188).

### 3.4. Chromosomal location of *bla*_CTX-M_ in the RefSeq assemblies

Among the 53 RefSeq environmental *E. coli* genomes, 20 carried *bla*_CTX-M_. Sixteen carried *bla*_CTX-M_ on plasmids and four carried *bla*_CTX-M_ on chromosomes (**Table S4**). None of the 20 RefSeq genomes with *bla*_CTX-M_ belonged to ST131, again indicating a potential difference in clonal composition of ESBLEC in environmental settings and clinical settings. The four genomes with chromosomal *bla*_CTX-M_ belonged to ST38 (n = 2), ST130 (n = 1) and ST648 (n = 1) and originated from various countries (Japan, New Zealand, Norway, and Switzerland), which indicates that diverse *E. coli* isolates carrying chromosomal *bla*_CTX-M_ are also present in environmental waters in other countries. In all four genomes, *bla*_CTX-M_ was associated with TPU ranging from 2,971 bp to 13,552 bp. A TPU containing multiple resistance genes besides *bla*_CTX-M_ and a TPU inserted into a multiresistance region were detected, again highlighting the importance of these mechanisms in the emergence of multidrug resistant *E. coli* in the environment (see **Figure S2** for the analysis of the genetic contexts of *bla*_CTX-M_ in these genomes).

ST38, which was detected in two RefSeq genomes containing chromosomal *bla*_CTX-M_, was also detected in two isolates with chromosomal *bla*_CTX-M_ sequenced in this study. Interestingly, ST38 was also found to be prevalent in some previous studies reporting chromosomal *bla*_CTX-M_ (Guenther et al., 2017; Guiral et al., 2011; Rodriguez et al., 2014). The reason for the prevalence of ST38 among *E. coli* with chromosomal *bla*_CTX-M_ is unknown. However, multiple integration events seem to have been involved in the acquisition of chromosomal *bla*_CTX-M_ in ST38 (see **Figure 1, Figure 2a**, and **Figure S2**), which potentially indicates ST38 has genetic mechanisms that can promote chromosomal integration of *bla*_CTX-M_.

### 3.5. Study limitations

This study has some limitations. The *E. coli* isolates analyzed in the present study were collected between 2011 and 2013 (i.e., ∼10 years ago). Additionally, a limited number of complete genomes were determined (n = 14), and the isolates were collected regionally, not nationwide. However, we also included RefSeq environmental *E. coli* genomes from various countries with various collection dates in our analysis, which might mitigate these limitations.

## 4. Conclusions

Here, we demonstrated that the genetic contexts of *bla*_CTX-M_ genes in environmental *E. coli* are highly diverse in terms of the associated mobile elements (various TPU, IS*CR1*, and IS*26*) and integration sites. This study also revealed that the chromosomal locations of *bla*_CTX-M_ genes were unexpectedly frequent in environmental ESBLEC in Japan. Chromosomal integration of *bla*_CTX-M_ might allow *E. coli* to stably maintain *bla*_CTX-M_ in environmental waters, though further studies are needed to confirm this.

## Acknowledgements

We acknowledge the NGS core facility of the Genome Information Research Center at the Research Institute for Microbial Diseases of Osaka University for the support in DNA sequencing. Computations were partially performed on the NIG supercomputer at ROIS National Institute of Genetics. This work was supported by the Environment Research and Technology Development Fund (JPMEERF20205006) of the Ministry of the Environment, Japan. This study was also supported in-part by the Program for the Development of Next-generation Leading Scientists with Global Insight (L-INSIGHT), sponsored by the Ministry of Education, Culture, Sports, Science and Technology (MEXT), Japan.

