## Supplementary Materials and methods, Supplementary Figures (S1-S2) for "Chromosomal integration of *bla*_CTX-M_ genes in diverse *Escherichia coli* isolates recovered from river water in Japan"

**Antibiotic susceptibility testing.**

Antibiotic susceptibility was assessed by microdilution using the dry plate Eiken assay (Eiken, Tokyo, Japan) using the antimicrobial agents listed in **Table S1**. The dry plate was designed to include antibiotic concentrations corresponding to both EUCAST epidemiological cutoff (ECOFF) values and CLSI breakpoints, where possible. We considered not only CLSI breakpoints (susceptible vs non-susceptible) but also ECOFF values (wild type vs non-wild type). This is because ECOFF values are epidemiologically based and thus are suitable for interpreting antibiotic susceptibility of environmental bacteria in comparison to clinical breakpoints, which are based on parameters relevant to therapeutic success. The dry plate was inoculated with an automated inoculation machine and incubated at 35 °C for 18 hours.

ESBL production was confirmed by disk diffusion using cefotaxime and ceftazidime disks with and without clavulanate (Eiken). The plate was incubated at 35 °C for 18 hours. A ≥ 5 mm increase in a zone diameter for either antimicrobial agent tested in combination with clavulanate compared with the zone diameter of the agent without clavulanate was considered ESBL-positive.

### Supplementary Figures

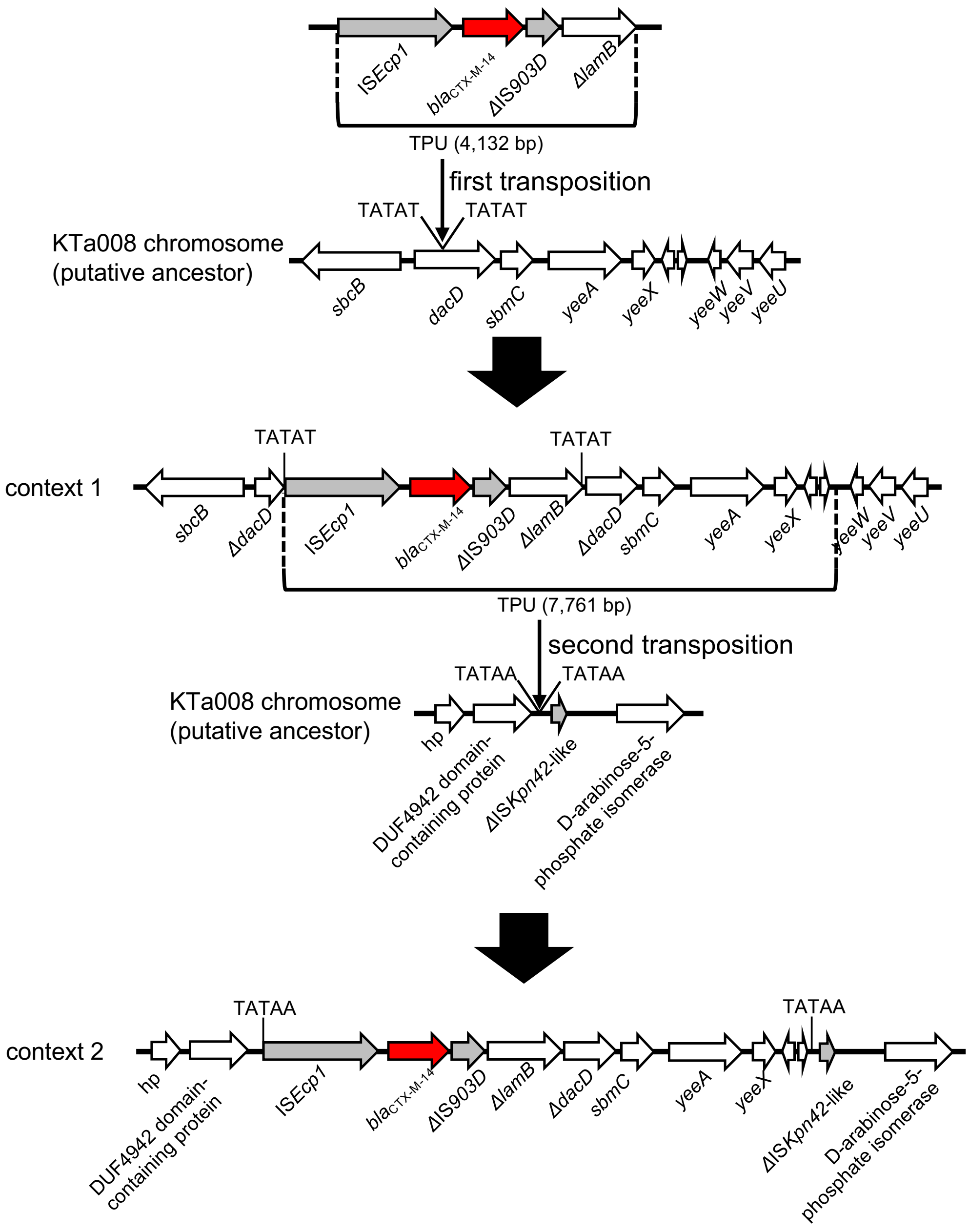


**Figure S1.** Suggested mechanism for generation of context 1 and context 2 in KTa008. First, a 4,132 bp TPU was inserted into *dacD,* generating TSD of TATAT. Next, a 7,761 bp TPU, containing the 4,132 bp TPU and an adjacent region, was transposed into another chromosomal region, generating TSD of TATAA. Red arrows indicate antibiotic resistance genes, gray arrows indicate mobile elements, and white arrows indicate other genes. hp: hypothetical protein.

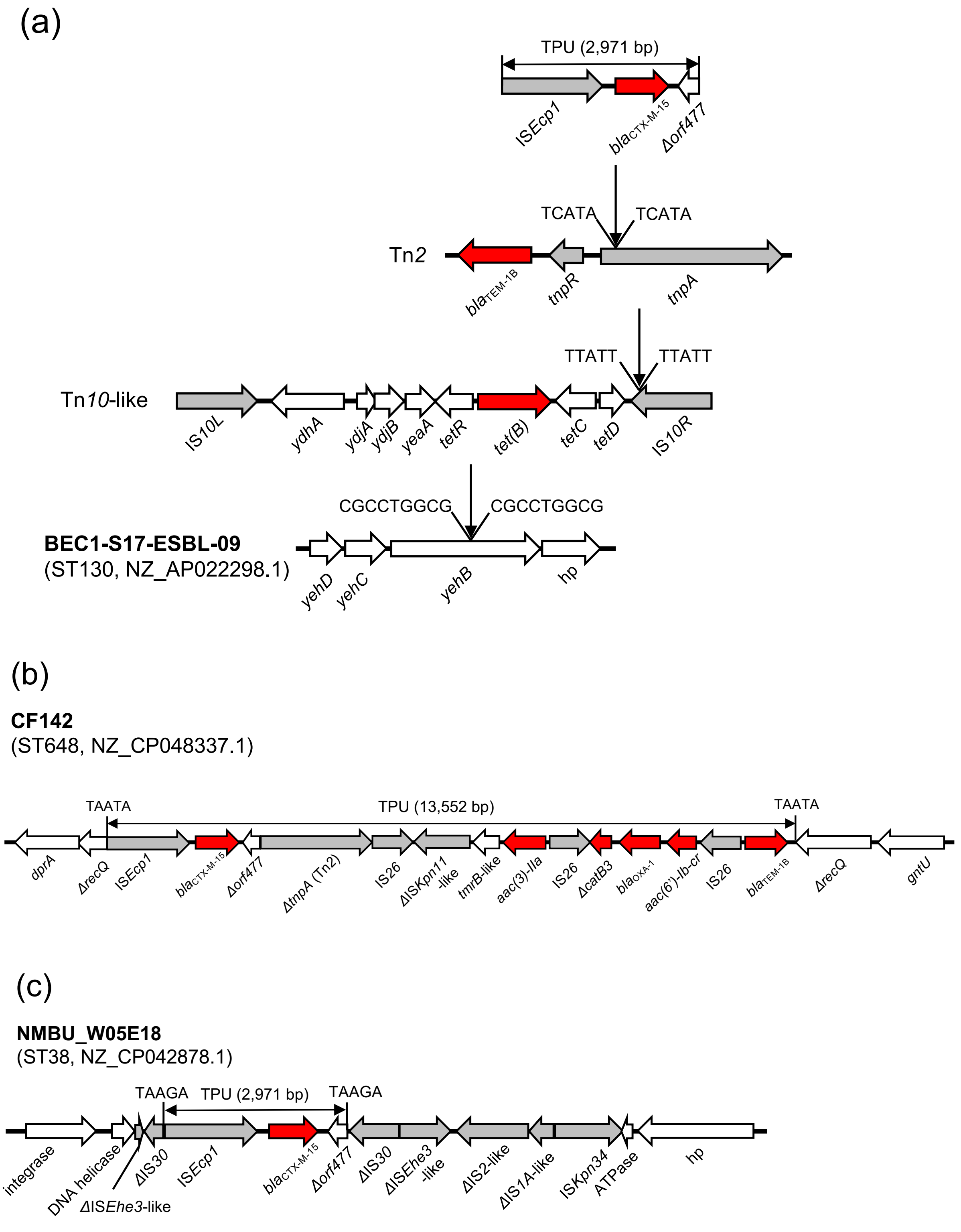
(a)
TPU (2,971 bp)
ISEcp1
blaCTX-M-15
Δorf477
TCATA
TCATA
Tn2
blaTEM-1B
tnpR
tnpA
TTATT
TTATT
Tn10-like
IS10L
ydhA
ydjA
ydjB
yeaA
tetR
tet(B)
tetC
tetD
IS10R
CGCCTGGCG
CGCCTGGCG
BEC1-S17-ESBL-09
(ST130, NZ_AP022298.1)
yehD
yehC
yehB
hp
(b)
CF142
(ST648, NZ_CP048337.1)
TAATA
TPU (13,552 bp)
TAATA
dprA
ΔrecQ
ISEcp1
blaCTX-M-15
Δorf477
ΔtnpA (Tn2)
IS26
ΔISKpn11 -like
tmrB-like
aac(3)-IIa
IS26
ΔcatB3
blaOXA-1
aac(6')-Ib-cr
IS26
blaTEM-1B
ΔrecQ
gntU
(c)
NMBU_W05E18
(ST38, NZ_CP042878.1)
TAAGA
TPU (2,971 bp)
TAAGA
integrase
DNA helicase
ΔISEhe3-like
ΔIS30
ISEcp1
blaCTX-M-15
Δorf477
ΔIS30
ΔISEhe3 -like
ΔIS2-like
ΔIS1A-like
ISKpn34
ATPase
hp

(d)

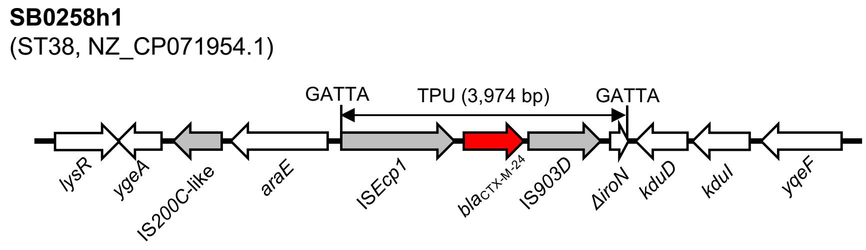


**Figure S2**. Genetic contexts of chromosomal *bla*$_{CTX-M}$ genes in RefSeq environmental *E. coli* genomes. (a) The genetic context of *bla*$_{CTX-M-15}$ in BEC1-S17-ESBL-09 (assembly accession no. GCF_014169915.1). A Tn*10*-like transposon was inserted into *yehB* in the chromosome, generating 9-bp TSD of CGCCTGGCG. Tn*2* was inserted in IS*10R* of the Tn*10*-like transposon, generating 5-bp TSD of TTATT. A 2,971 bp TPU containing *bla*$_{CTX-M-15}$ was inserted into *tnpA* of Tn*2*, generating 5-bp TSD of TCATA. (b) The genetic context of *bla*$_{CTX-M-15}$ in CF142 (assembly accession no. GCF_010365385.1). A 13,552 bp TPU, which carries *bla*$_{CTX-M-15}$ and other resistance genes, was inserted in *recQ* in the chromosome, creating 5-bp TSD of TAATA. (c) The genetic context of *bla*$_{CTX-M-15}$ in NMBU_W05E18 (assembly accession no. GCF_013282275.1). A 2,971 bp TPU carrying *bla*$_{CTX-M-15}$ was inserted in IS*30*, generating 5-bp TSD of TAAGA. The TPU was surrounded by multiple ISs. (d) The genetic context of *bla*$_{CTX-M-24}$ in SB0258h1 (assembly accession no. GCF_017603565.1). A 3,974 bp TPU carrying *bla*$_{CTX-M-24}$ was inserted in an intergenic region in the chromosome, generating 5-bp TSD of GATTA. The references for the four genomes are as follows: BEC1-S17-ESBL-09 (Sekizuka et al. *Research Square*. 2020. DOI:10.21203/rs.3.rs-52275/v1.), CF142 (Bleichenbacher et al.

*Environmental Pollution*. 2020. DOI: 10.1016/j.envpol.2020.115081.), NMBU_W05E18 (Finton et al. *Frontiers in Microbiology*. 2020. DOI: 10.3389/fmicb.2020.01273.), and SB0258h1 (Gray et al. *Microbiology Resource Announcements*. 2021. DOI: 10.1128/MRA.00328-21.).

**Supplementary Tables**

See the Excel file for the following tables:

**Table S1**. Basic characteristics of 18 ESBLEC isolates.

**Table S2**. Basic characteristics of each replicon of ESBLEC genomes sequenced to closure in this study.

**Table S3**. Results of BLASTN searches of TPU and the surrounding regions against the nr/nt database.

**Table S4**. Basic characteristics of RefSeq environmental *E. coli* genomes with $bla_{\text{CTX-M}}$.
